# Automatic sleep stage classification using physiological signals acquired by Dreem headband

**DOI:** 10.1101/2023.10.05.561041

**Authors:** Shahla Bakian Dogaheh, Mohammad Hassan Moradi

## Abstract

In this paper, we aim to propose a model for automatic sleep stage classification based on physiological signals acquired by Dreem Headband and extreme gradient boosting (XGBoost) method. The dataset used in this study belongs to a challenge competition, namely as “Challenge Data”, held in 2017-2018, and is publicly available on their website. Recordings, includes 4 EEG channels (FpZ-O1, FpZ-O2, FpZ-F7, F8-F7), 2 Pulse oximeter (RED & infra-red), and 3 accelerometer channels (X, Y, Z). In this work, sleep stages have been scored according to the AASM standard. Different features were extracted from the physiological signals after applying a preprocessing step. Each of the elicited features from EEG and PPG signals is falling into one of the three categories: time-domain, frequency domain, or non-linear features. Moreover, ancillary features including body movement, frequency features, breathing frequency, and respiration rate variability were also extracted from the accelerometer signal. Significance of the extracted features was examined through the Kruskal Wallis test, and features with P-value>0.01 were discarded from features set. Finally, significant features were classified by using support vector machine (SVM), K-nearest neighbors (KNN), random forest (RF), and XGBoost classifiers. Due to the class imbalance problem, repeated stratified 5-fold cross-validation was performed in order to tune systems parameters. Results show that among the four above-mentioned models, XGBoost has the best performance for the 5-class classification problem with accuracy: 81.34%±0.76% and Kappa 0.7388±0.0101. The proposed model shows promising results, therefore the model can be implemented in Dreem headband to differentiate between sleep states efficiently and be applicable in clinical trial.

## 1. INTRODUCTION

Sleep is a physiological state that is characterized by reversible loss of consciousness and changes in respiration, brain wave activity, and other physiological functions. Sleep accounts for one-third of humans’ lifespan and plays an important role in physical and mental health. Studies show that poor quality sleep and sleep disorders not only increase the risk of depression, diabetes, neurological and cardiovascular disease but also adversely affect cognitive function such as learning, attention, and memory in the long run [1],[2],[3]. Nowadays, according to the International Classification of Sleep Disorder (ICSD-II) criteria, 84 different sleep disorders have been identified [4]. Diagnosing and treating sleep disorders relies on detecting different sleep states accurately; therefore, sleep stage scoring is essential. To this end, a set of biomedical signals including electroencephalogram (EEG), electromyogram (EMG), electrooculogram (EOG), and electrocardiogram (ECG), called polysomnography (PSG) recordings, are acquired [5]. Among these signals, EEG signal is regarded as an important diagnostic tool in sleep medicine [6]. PSG recordings are then segmented into 30-second epochs and scored by sleep experts according to one of the sleep scoring guidelines, Rashthaffen and Kilz (R & K) [7] or American Academy of Sleep Medicine (AASM) [8].

Manual sleep scoring is deemed to be a subjective, time consuming, and skill dependent task that can bring about disagreement about sleep scoring among specialists and result in misdiagnosis of sleep disorders. Therefore, developing an automatic sleep staging model may address this problem.

To date, Various studies have been conducted for this purpose and designed sleep stage identification systems based on using different signals [9], [10] features [11], [12] and classifiers [13],[14], [15], [16]. These methods rely on either single-channel processing, which is based on using single-channel EEG recording [17], [18], [19], or multi-channel processing [20]. In the latter approach, the PSG recordings are employed for extracting informative features. Although using a combination of multi-channel signals result in higher performance, this approach is less desirable since recording equipment disrupt patient’s sleep.

The next step in designing an automatic sleep scoring system is extracting informative features from physiological signals. In general, extracted features can be divided in to several categories including time domain [19], [21], [22], frequency domain [23], [24], [25], time-frequency feature [26], nonlinear features [27], [28] and features based on using convolution neural networks [29], [5], [30]. After extracting discriminative information from signals, various classifiers [31], [24] can be utilized to assign a sleep state to each epoch. In this regard, some of the proposed systems is presented here. Supratak et al. used 62 healthy subjects’ recordings from MASS and Sleep-EDF datasets and proposed an end-to-end automatic sleep stage classification model based on single-channel EEG signal and a deep learning model. Their model’s accuracy for 5-class classification problem is 86.2% and 78.9% on MASS and Sleep-EDF datasets, respectively [5]. In another study, Zhang et al. proposed new time-domain features using a sequence merging method in pre-processing step. They evaluate their model on a data recorded from three participants with breathing-related sleep disorder[32]. Tripathy et al. decomposed RR-times series into intrinsic mode function (IMF), and extracted features from IMF of RR-time series and EEG signal to distinguish between different sleep stages. Their model achieved the following accuracy: for wake vs sleep: 85.51%, light sleep vs deep sleep: 94.03%, NREM vs REM stages: 95.71% [33]. Aboalayon et al. used the support vector machine (SVM) classifier and 13 subject’s data to perform sleep stage classification based on single-channel EEG. The accuracy obtained for classifying five classes was 92.5% [34]. In some other studies, the viability of using wearable sensors’ data in sleep stage classification was assessed. For instance, Aktaruzzaman et al., used 18 healthy subjects’ data and investigate the possibility of performing sleep stage scoring using ECG, wrist and chest actigraphy signals acquired by wearable sensors [1]. Beattie et al. used 60 healthy subjects’ data acquired from a wrist-worn device and investigated the possibility of 4-class sleep stage scoring using accelerometer and Photoplethysmography (PPG) signals. The accuracy and kappa for 4-class was 69% and 0.52±0.14, respectively [35]. In another study, Yuda [36] used heart rate and actigraphy signals to distinguish REM stage from Wake stage and NREM stage from other stages. In this study, 289 subjects, consisting 40643 epochs, participated. Their model’s accuracy for classifying REM vs. other classes and REM vs. Wake stage was 75.8% and 74.5%, respectively.

Even though various automated models have been presented, there are shortcomings with a large number of proposed models in practice that make them inapplicable for clinical trial [37], [18]. Therefore, there is still a need for accurate, robust, and portable models, which are capable of recording large datasets as well as allowing reasonable evaluation of patient signals [38]. Because one of the challenges in this context is that proposed models should have a promising performance on patient data. However, most of the studies only utilize healthy subjects’ recordings in their models, so the results can’t be reliable. Another challenge that arises in this setting is the traditional way of recording polysomnography data. Some of these limitations are listed as follows:

1. The recording process is complicated and time-consuming.
2. A trained sleep specialist is needed for recording data
3. It is expensive and costly.
4. Besides the stressfulness of Clinical environments and devices, recording equipment is cumbersome and inconvenient to patients.
5. The recorded signals are not very reliable to indicate a person’s normal sleep and can lead to inaccurate sleep disorder diagnosis since polysomnography is usually recorded at one night.

While PSG is the gold standard for sleep assessment, in recent years, there is an increasing interest in developing new groups of wearable devices (e.g., headbands, bracelets, smartwatches, or chest actigraphs) that provide the opportunity for home sleep-monitoring [35], [39]. These devices can record EEG, PPG, heart rate, respiration, body movement, and more signals based on which sleep assessment can be carried out in a real-world environment compared to a sleep laboratory [40]. Furthermore, unlike traditional PSG, these wearable devices are portable, smaller in size, cheaper, and less obtrusive.

In this study, we used Dream headband’s signals as an alternative to PSG recordings to overcome the limitation of multi-channel processing and leveraged machine learning method to propose an appropriate automatic sleep scoring model. This paper is an extended version of our previous work published in [41]. The paper organization is as follows: in section 2, the proposed methodology, which includes dataset, extracted features, statistical test, classifiers and stratified K-fold cross-validation are cited. In sections 3, 4, and 5 results, discussion, and conclusion are discussed, respectively.

## 2. METHOD

State-of-the-art automatic approaches for sleep stage classification are divided into two categories depending on whether a convolutional neural network (CNN) is being used to learn the features from raw data or they are extracted using expert knowledge [42]. In this paper, an automatic sleep scoring system based on extracting handcraft-features from physiological signals, the latter approach, is proposed. The XGBoost model and three broadly known machine learning methods are adopted in order to identify five sleep states according to AASM guidelines. Figure 1, shows the block diagram of proposed model for 5-class sleep stage classification.

**Figure 1.**
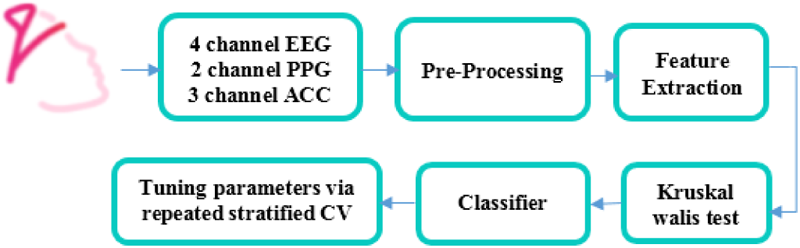
Block diagram of proposed method.

### 2.1. Dataset

A publicly available sleep dataset that was used in Challengedata’s challenge [43], held at 2017-2018, has been employed in this study. The dataset was recorded by Dreem headband (DH) [44], Figure2, and the recording includes four EEG channels (Fpz-F7, Fpz-O2, FPZ-O8, and F8-F7) sampled at 125Hz, two pulse oximeter channels (red-infrared) sampled at 50Hz, and a 3-D accelerometer signal sampled at 50Hz. The dataset contains training and test sets, which involve 43830 epochs (850 subjects) and 20592 epochs (879 subjects), respectively. Each 30-s epoch of the signals was manually scored as wake, NREM1, NREM2, NREM3, or REM stage according to the AASM criteria. The number of epochs available for each sleep stage in the training set, and the number of epochs used in this study are given In Table 1. Herein, the proposed model is trained using training set in the challenge dataset. The data is freely available through challenge website at [43].

**Figure 2.**
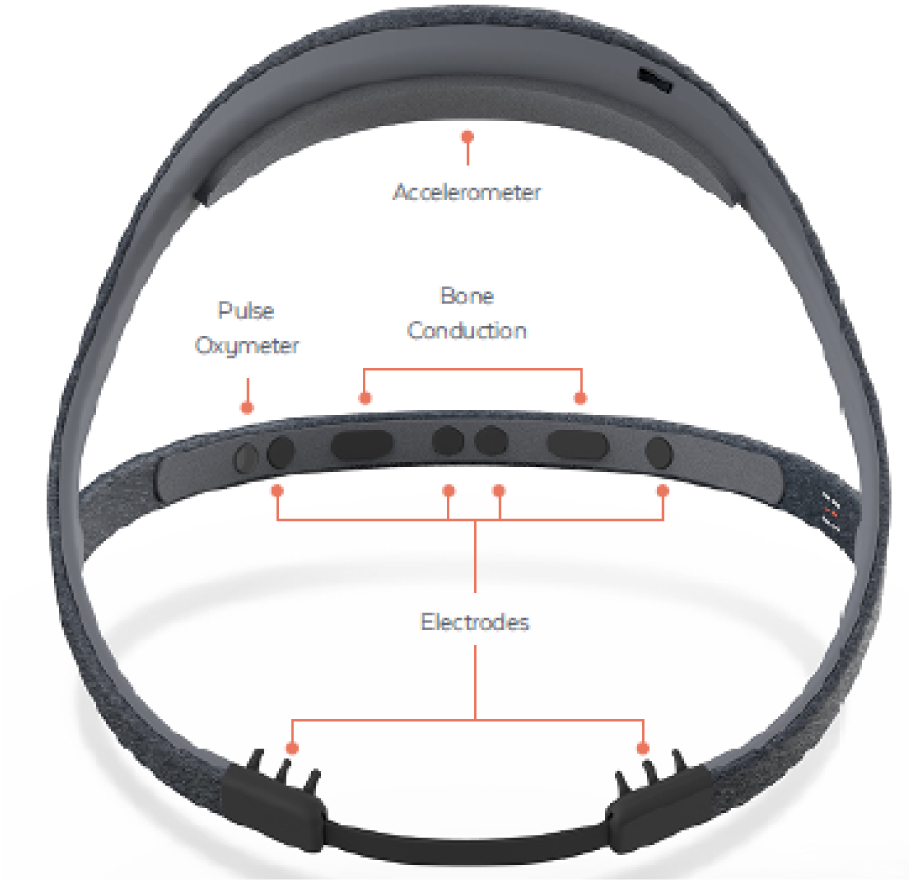
Dreem headband [44].

**Table 1.** The number of epochs available for sleep stages in dataset and used in the study.

| Sleep stages | Number of epochs in the dataset | Number of epochs used |
| --- | --- | --- |
| Wake | 4939 | 2470 |
| NREM1 | 1359 | 1359 |
| NREM2 | 16139 | 8070 |
| NREM3 | 13780 | 6890 |
| REM | 7613 | 3807 |
| Total | 43830 | 22596 |

### 2.2. Pre-processing

In order to increase the signals quality a preprocessing step is applied to signals before feature extraction. Accordingly, the signals are filtered as follow:

#### EEG signals

The EEG signal is filtered with a 14th order Butterworth bandpass filter within the frequency range of 0.5 to 18 Hz.

#### Pulse oximeter signals

The pulse oximeter signal is filtered using a 4th order Butterworth low pass filter with a cut-off frequency of 3 Hz. Then in order to have a smooth signal exponential smoothing method is applied with alpha 0.3.

#### Accelerometer signals

The accelerometer signal is filtered using a 4th order Butterworth bandpass filter within the frequency range of 0.16 to 0.3 Hz.

### 2.3. Feature extraction

Various features are extracted from EEG, PPG, and accelerometer signals due to their potential in distinguishing between the characteristics of sleep stages. In this section, a brief description of these features are presented.

#### 2.3.1. EEG signal

In general, extracted features from the EEG signals are divided into three main categories containing time domain, frequency domain, and nonlinear features. According to the AASM guideline, sleep stages can be identified through changes in amplitude and frequency band of EEG rhythms. Therefore, time domain and frequency domain features are suitable in determining sleep stages. Although fractal-based and entropy features cannot show these changes, they are beneficial in detecting sleep stage transitions [2].

##### Time domain features

In general, according to the AASM guideline, the amplitude level of EEG signal is an important feature that can distinguish between different sleep stages. Therefore, maximum and median of EEG amplitude are extracted.

##### Maximum

The largest value in time series.

##### Median

The middle value of ordered numbers.

##### Number of zero crossing (NZC)

The number of times a signal crosses the baseline is called number of zero crossing. It is an important feature in sleep staging since it can extract information about specific waveforms, e.g. sleep spindles which have a higher frequency and a higher number of zero crossing.

##### Standard deviation (SD)

The standard deviation measures the dispersion of data and for a given signal of x(n), it is calculated as follow:

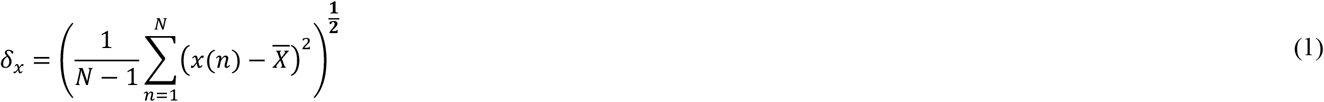

##### Skewness

Skewness quantifies the asymmetry behavior of a signal. For a given signal x(n) with N sample x_i_, Skewness can be defined as follows:

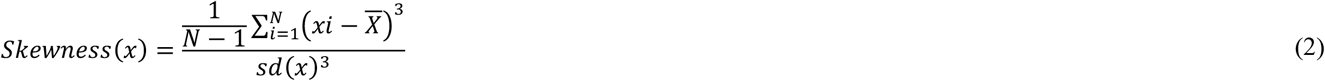

##### Kurtosis

Kurtosis is a measure of the tailedness of a probability distribution and defines as follows:

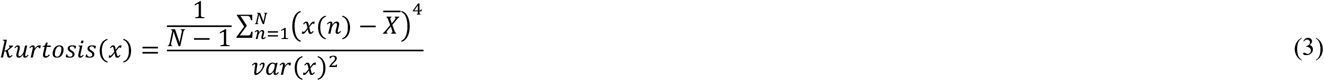

##### Hjorth parameters

Hjorth mobility and complexity are a good estimate of mean frequency and spectral bandwidth, and it can extract information about the stage-specific waveform. These parameters are defined as follow:

Hjorth mobility (HM):

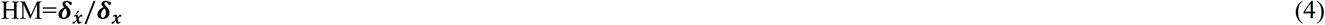

Hjorth complexity (HC):

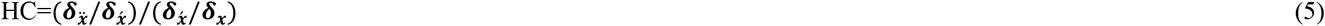

Where 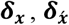 and 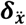 are standard deviation of x(n), the first and second derivatives of x(n) (x′(n) and x″(n)), respectively

##### Frequency domain features

According to the AASM rules, sleep stages can be distinguished from one another by the presence of EEG rhythms (e.g., alpha, delta, theta, and beta wave) in each state. Thus, the power spectra of each EEG rhythm can provide valuable information about sleep stages. To this end, after windowing the EEG signal to non-overlapping 125 sample length segments, power spectral density (PSD) and power ratio of each EEG rhythm are computed by means of Welch spectral density estimation. The EEG waveforms and frequency domain features are represented in Table 2 and Table 3, respectively.

**Table 2.** Frequency band of EEG signal.

| EEG Waveforms | $\delta$ | $\Theta$ | $\alpha$ | $\sigma$ |
| --- | --- | --- | --- | --- |
| Frequency band | 0.5-4 Hz | 4-8 Hz | 8-12 Hz | 12-15 Hz |

**Table 3.** Frequency Features.

| PSD | Power ratios: |
| --- | --- |
| Delta band power ($P_\delta$) | $\frac{P_\delta}{P_\theta}$ |
| Theta band power ($P_\theta$) | $\frac{P_\delta}{P_\alpha}$ |
| alpha band power ($P_\alpha$) | $\frac{P_\delta}{P_\sigma}$ |
| sigma band power ($P_\sigma$) | $\frac{P_\theta}{P_\alpha}$ |
| | $\frac{P_\theta}{P_\sigma}$ |
| | $\frac{P_\alpha}{P_\sigma}$ |

**Table 4.** Performance metrics of XGBoost model for 5 class classification problem.

| Confusion Matrix and Statistics |  |  |  |  |  |
| --- | --- | --- | --- | --- | --- |
|  | Wake | NREM1 | NREM2 | NREM3 | REM |
| Wake | 294 | 14 | 26 | 3 | 15 |
| NREM1 | 8 | 87 | 15 | 0 | 13 |
| NREM2 | 46 | 74 | 953 | 65 | 147 |
| NREM3 | 11 | 4 | 62 | 961 | 3 |
| REM | 17 | 13 | 68 | 2 | 399 |
| Statistics by Class: |  |  |  |  |  |
| Sensitivity | 0.78191 | 0.45312 | 0.8479 | 0.9321 | 0.6915 |
| Specificity | 0.98016 | 0.98842 | 0.8474 | 0.9647 | 0.9633 |
| Overall Statistics |  |  |  |  |  |
| Accuracy | 0.8164 |  |  |  |  |
| Kappa | 0.7478 |  |  |  |  |

##### Non-linear feature

EEG signal is non-stationary in nature and behaves irregularly. Entropy and fractal-based features can be a good measure of irregularity and complexity in this signal because the signal irregularity is directly related to the information content, entropy, and fractal values. During sleep, changes in sleep stages result in changes in the amplitude and frequency content of the EEG signal. These changes can be finely represented by entropy and fractal-based features [45], [2]. Herein, nonlinear features extracted from EEG signal include Rényi entropy, Relative spectral entropy, Katz and Higuchi fractal dimension. These features are defined as follows:

##### Rényi entropy (RE)

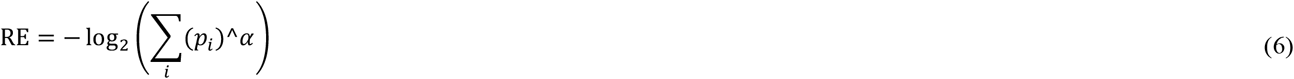

Where *p* is probability distribution and α is the order of Rényi entropy, in this paper α =2.

##### Relative spectral entropy

Relative spectral entropy of a signal is a measure of its spectral power distribution

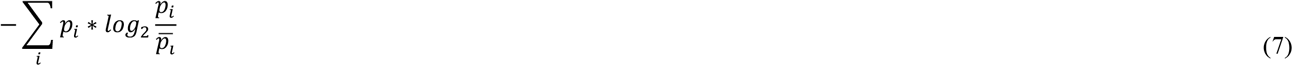

P = normalized power spectrum

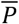 = normalized mean power spectrum

##### Katz Fractal Dimension (KFD)

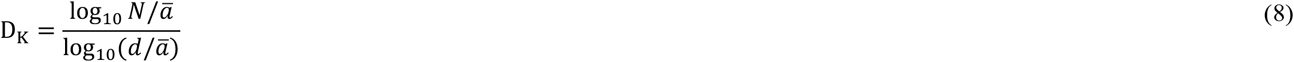

**N:** is the total length of the curve or sum of distances between successive points

**d:** the distance between the first point of the sequence and the point of the sequence that provides the farthest distance

ā: Normalizing distances between successive points

The KFD’s formula and description is Adapted from [46].

##### Higuchi’s Fractal Dimension (HFD)

HFD is extracted due to its capability for detecting the transition between NREM and REM states and its ability in demonstrating specific variation of sleep EEG across sleep cycle. Details for computing HFD is described in [47].

#### 2.3.2. Pulse oximeter

From Pulse oximeter signal, sample entropy (SE) and statistical features, including mean, max, median, and standard deviation, are extracted [48] using Red and Infra-Red channels.

Heart Rate (HR) have also been extracted from the Red channel. But after applying the statistical test, no significant difference among sleep stages was shown, which can be because of deficient resolution of the signal; thus, HR was discarded from feature set.

#### 2.3.3. Accelerometer

Four informative features, including body movement, frequency features, breathing frequency, and respiration rate variability (RRV), are extracted from the accelerometer signal.

##### Body movement

In order to track general motion independent of direction, E(t) is calculated using the Euclidean sum formula, thus it will be less sensitive to patients’ head position. E(t) define as:

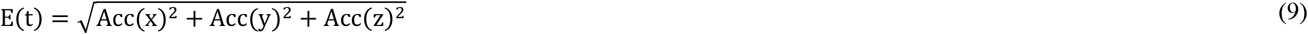

Where ACC(x), ACC(y), ACC(z) refer to three accelerometer channels X,Y, Z, respectively. Then mean, median and max of upper and lower envelopes of E(t) signal is calculated.

These features can offer information about body movement, which will be useful in detecting Wake stage.

##### Frequency features

In order to extract information about breathing and body movement, the following steps had been taken:

Step1: calculating Instantaneous Frequency (IF) of 1st principal component of three accelerometer channels (x, y, z). To this end Hilbert [49] method was used for calculating IF.

Step2: calculating the mean, median, and standard deviation of the first step’s output.

We believe that extracting the first principle component of three accelerometer channels can have information about breathing and instantaneous frequency can mark irregular breathing. These information can be useful in distinguishing REM states from non-REM [50] as breathing is irregular during REM state.

##### Breathing frequency

To extract breathing frequency, a similar method as presented in [51] is used. To this end, Z-axis of the accelerometer signal has been utilize and the following steps have been taken to calculate it.

1. The signal is filtered with a Butterworth band-pass filter between 0.16-0.3 Hz, then zero-crossing method is applied in order to compute mean breathing frequency (f_s_).

2. The signal is again filtered between 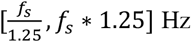, and the minimum (fs_min_) and maximum (fs_max_) breathing frequency are computed after applying zero-crossing method.

3. Finally, the signal is filtered between [fs_min_-fs_max_] and after applying zero crossing method, breathing frequency is obtained [51].

##### Respiration Rate Variability (RRV)

The method employed for computing RRV feature is the same method presented in [51], [52], and contains four steps:

Step1: All positive values are eliminated from the accelerometer signal to keep the expiratory part.

Step2: Fast Fourier transform of the expiratory part of the signal is computed, then H1/DC ratio is measured from its spectrum, where H1 and DC are defined as:

H1: Fundamental frequency’s amplitude (In this paper, the first peak after zero frequency is considered as H1.) DC: The zero-frequency in the spectrum of the expiratory signal.

Step3: Then RRV is obtained as follow:

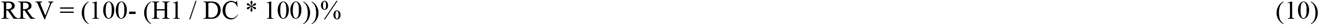

Step 4: Due to sensor displacement, RRV> 85% are discarded, and nearest neighbor interpolation method was employed to replace the removed points [52].

It should also be noted that after computing RRV value for each accelerometer’s channel, the minimum value is selected to reduce the noise. [51].

### 2.4. Statistical test

The statistical significance of each extracted feature is investigated by Kruskal–Wallis test, which is a non-parametric equivalent of the one-way ANOVA. In order to find a discriminative feature set that can distinguish between different sleep stages, features with p-values<0.01 are considered, and the rest are discarded from the feature vector. After applying Kruskal–Wallis test p-value of all features except Heart rate were <0.01 and ensure that selected features statistically differentiate between sleep stages and, therefore, can be suitable for automatic classification of sleep stages.

### 2.5. Classification

In this study, a prediction model based on the XGBoost classifier [53] is proposed to classify the elicited features. XGBoost is an ensemble machine learning algorithm that converts a weak classifier into a strong learner. XGBoost is based on efficient implementation of gradient boosted trees algorithm, which is an iterative technique that adjusts the weight of observation based on the information from a previously grown tree one after the other and tries to improve its prediction in the subsequent iteration. The benefits of using XGBoost algorithm are that it is ten times faster than the existing algorithms, scalable in all scenarios, works well for unbalanced datasets, and is known for its high predictive performance [42],[43]. Mathematically, the model can be represented as follows:

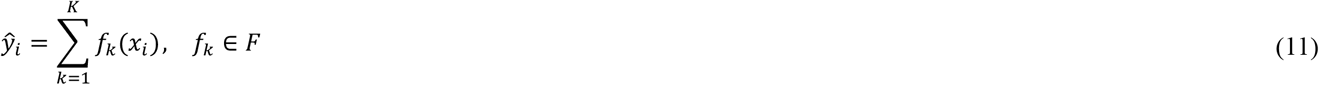

Where ŷ is the prediction value, *x*_*i*_ is the feature vector, K is the number of trees, *f* is a function that includes tree structure and the leaf scores, and F is the set of all classification and regression trees (CART). The algorithm is trained by iteratively adding a tree in each iteration to increase the model’s performance. Similarly, the prediction value at the t step is defined as:

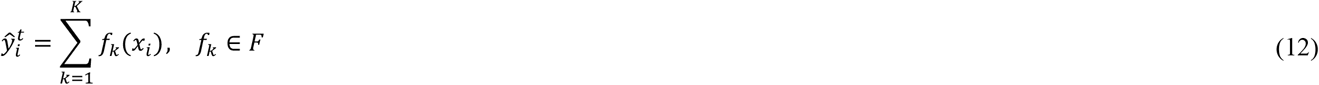

In order to train the model, an objective function should be optimized. The objective function measures how well the model fits the training data and controls the simplicity. It is defined as follows:

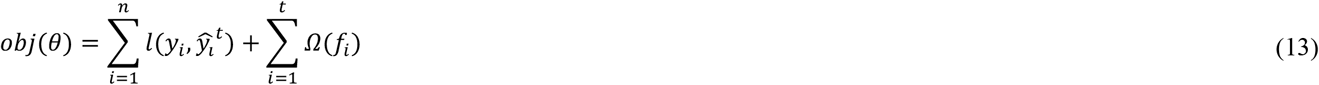

Where the first term is the training loss function and the second term is the regularization term, which controls the complexity of the model and prevents overfitting.

The regularization term is also defined as:

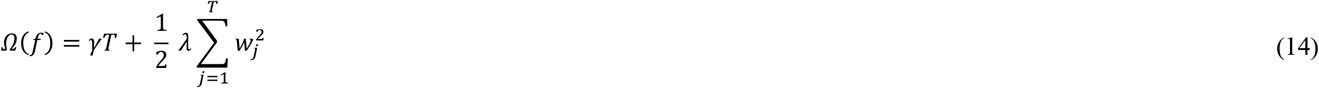

Where T is the number of leaves, 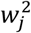 is the score on the j-th leaf, and *γ* and *λ* control the regularization. For learning the function*f*_*i*_, the objective function (Eq. 15) should be optimized.

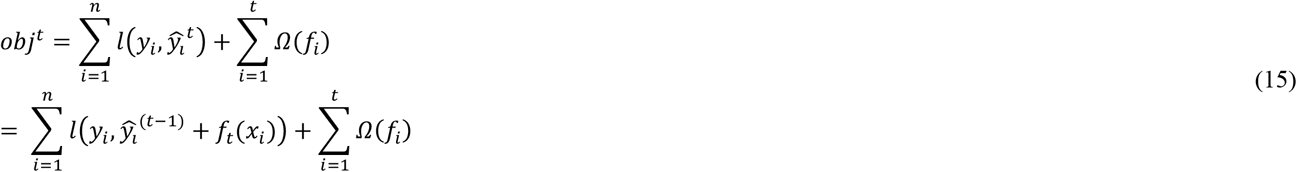

Since traditional optimization cannot be used, second order Taylor approximation is utilized:

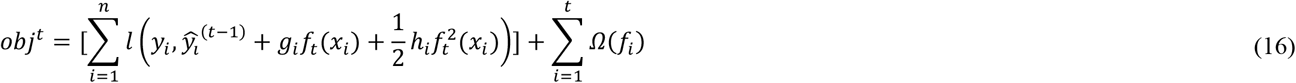

Where *g*_*i*_ and *h*_*i*_ are:

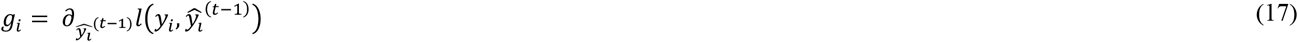

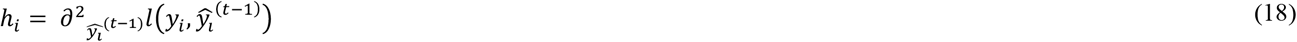

Each tree can be defined as:

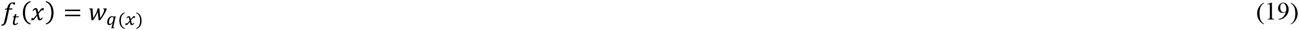

Where q is a function assigning each data point to the corresponding leaf, w is the vector of scores on the leaves and T is the number of leaves.

So the objective function at the t-th step can be written as follows:

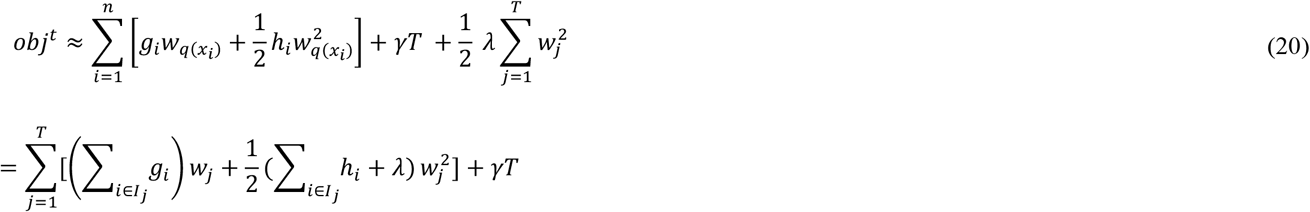

Where the set *I*_*j*_ = { *i*|*q*(*x*_*i*_) = *j*} includes the indices of data points assigned to j-th leaf.

In order to optimize the objective function, the quadratic problem of Eq. (20) should be solved. The leaf scores can be calculated as follows:

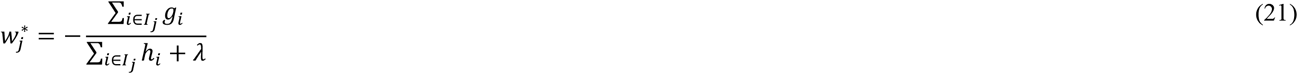

Finally the optimal value can be calculated as:

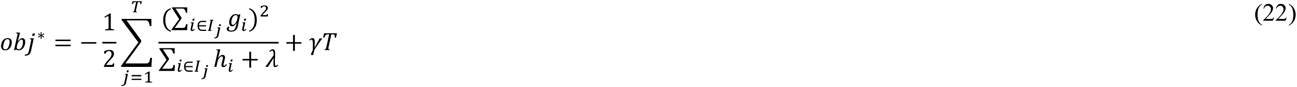

[53].

XGBoost also offers a range of hyperparameters to tune the behavior of the algorithm. These parameters are divided into three categories: booster parameters, general parameters, and learning task parameters. Herein, multi logarithmic loss function was used as an input for the objective function to be optimized, and the maximum number of trees was set to 2500. The early stopping rounds is set to 20, therefore training with the validation set will stop if the performance didn’t improve for 20 rounds. In order to tune the model’s hyperparameters to achieve the highest possible performance, we performed grid searches and repeated stratified 5-fold cross-validation with 10 times repeat for each set of hyperparameters. The tuned hyperparameters are as follow: the learning rate is set to 0.1, the minimum weight of a child tree is set to 1, gamma is set to 9.15, the subsample ratio of the training instance is set to 1, the subsample ratio of columns when constructing each tree is set to 0.5, and maximum depth of the tree is set to 7.

In this study, we also draw a comparison between XGBoost classifier and three other supervised machine learning algorithms including Random forest (RF), support vector machine (SVM), and k-nearest neighbours (KNN), in terms of their performance and runtime. The ML methods were implemented using R packages. We also investigate the effectiveness of adding each signal on classification performance. The results are mentioned in section 3.

### 2.6. Evaluation

In classification process, the whole dataset were randomly split into training and test sets, in an 85:15 ratio, respectively. For tuning models’ hyperparameters, repeated stratified K-fold cross-validation (CV), with K=5 and n=10 times repeat, is performed on the training set. Then the test set was used to evaluate the performance of the proposed model. One of the benefits of this approach is that it can make up for CV’s drawback. Strictly speaking, K-fold cross-validation split the whole dataset into K folds, in which one of the fold is the test set and the rest is considered as the training set. The most significant drawback of this approach is that in unbalance datasets, where the number of samples participating in each class is not equal, by random sampling, some classes may not participate in sampling or maybe very few, so the model is not well trained. But stratified CV spilled each class into K folds and use one part for test and the rest for training; therefore, it can hold the same ratio of classes across each CV split. In this study, the Accuracy, Cohen’s Kappa, specificity and sensitivity measures were utilized to evaluate the proposed method’s performance. Definitions of these measures are as follows:

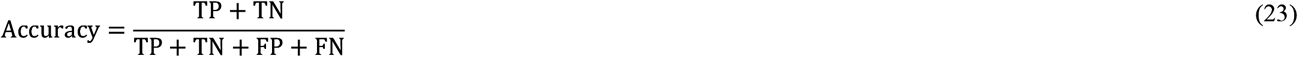

Where TP, TN, FP, and FN refer to number of true positive, true negative, false positive and false negative.

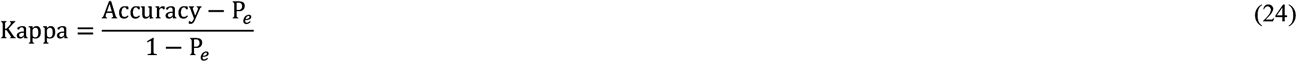

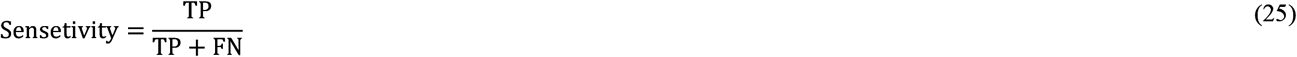

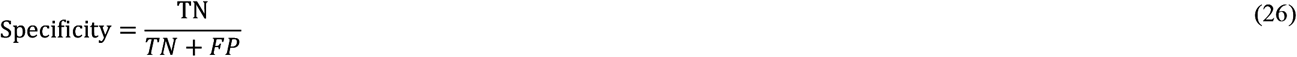

The accuracy index and Cohen’s Kappa coefficient are used to evaluate overall sleep stage classification performance, and specificity and sensitivity measures are used to evaluate each sleep stage classification performance.

## 3. RESULTS

This study endeavors to propose a functional automatic sleep staging system using Dreem headband’s signal, which is a wireless FDA-registered medical device that can measure sleep activity [44],[54]. To this end, the dataset obtained from ChallengeData’s challenge competition and after a pre-processing step, a set of features, including: time domain, frequency domain, non-linear domain, body movement, breathing frequency, and respiration rate variability are extracted from 4 EEG, 2 pulse oximeter, and 3 accelerometers channels. The extracted features can represent the information content of signals, and the Kruskal Wallis test examined it. To perform classification XGBoost, RF, SVM, and KNN classifiers were applied and the influence of adding each signal’s feature on the system’s performance was also investigated for all classifiers. The results are listed in Figure 3, 4. As shown in Figure 3, 4, among the compared classifiers, XGBoost model outperforms other classifiers (In the presence of all features) in terms of accuracy & kappa with a mean accuracy of 81.34± 0.76 and kappa 0.7388±0.0101. Although the XGBoost and RF performance metrics are not much different, XGBoost’s runtime is far less than RF’s and other classifiers, which shows this model is more efficient.

**Figure 3.**
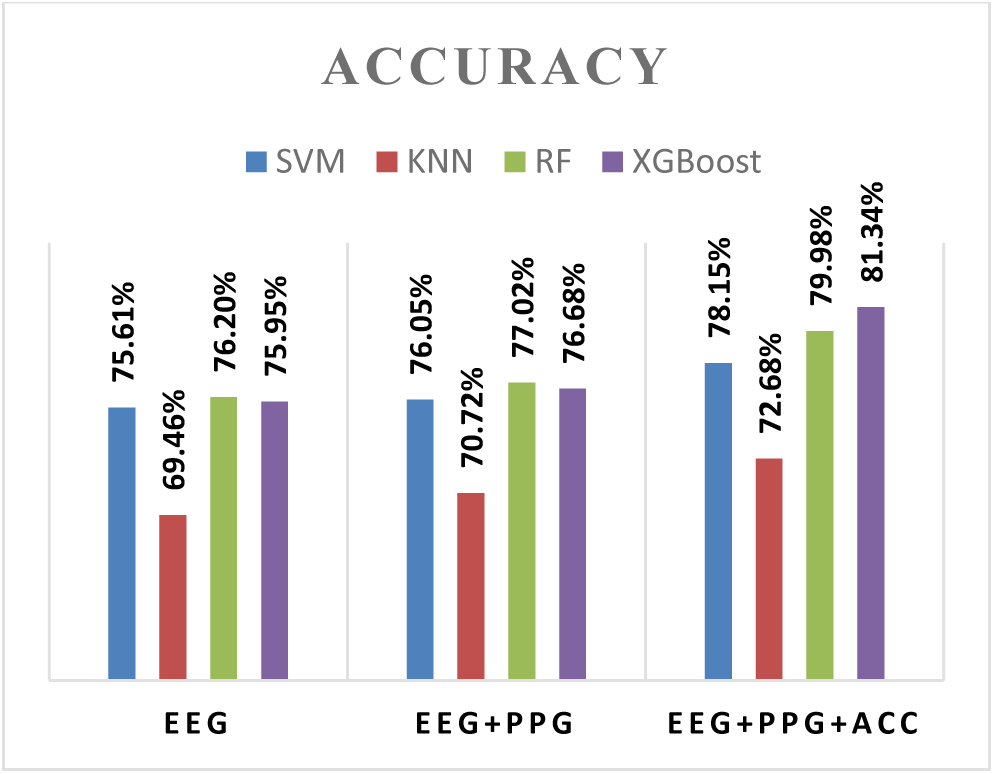
Comparison of different classifiers in term of accuracy and investigating each signal effectiveness.

**Figure 4.**
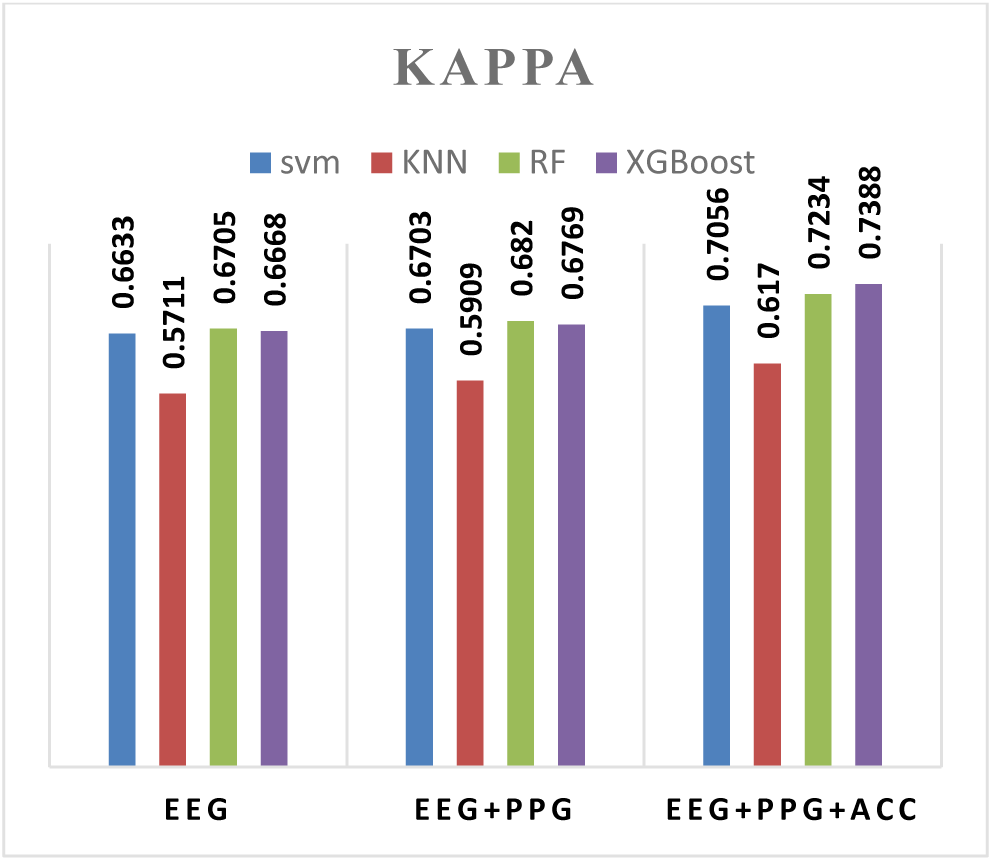
Comparison of different classifiers in term of kappa and investigating each signal’s effectiveness.

One of the key factor in sleep stages scoring is to propose a model that can identify different sleep stages that is crucial in diagnosing sleep-related disorders. For instance, for diagnosing sleep apnea, it is vital to differentiate sleep and wake states accurately. Therefore, different sleep stage classification are done to address this problem, and the model’s performance in terms of accuracy, Cohen’s Kappa, specificity and sensitivity are given in Table 5, 6. For 2-class classification problem (Wake vs Sleep) the model’s accuracy is 96.15% with 95% confidence interval (CI): (0.9544, 0.9678) and Kappa: 0.7857. For 3-class classification problem that distinguish Wake, NREM and REM stages the proposed model obtained 88.15% accuracy (95% CI: (0.87, 0.8924)) and kappa: 0.7154.

**Table 5.** Performance metrics of XGBoost model for 3-class classification problem.

| Confusion Matrix and Statistics |  |  |  |
| --- | --- | --- | --- |
|  | 3 Class |  |  |
|  | Wake | NREM | REM |
| Wake | 274 | 29 | 13 |
| NREM | 86 | 2261 | 170 |
| REM | 7 | 86 | 374 |
| Statistics by Class: |  |  |  |
| Sensitivity | 0.74659 | 0.9516 | 0.6715 |
| Specificity | 0.98568 | 0.7229 | 0.9661 |
| Overall Statistics |  |  |  |
| Accuracy | 0.8815 |  |  |
| 95% CI | (0.87, 0.8924) |  |  |
| Kappa | 0.7154 |  |  |

**Table 6.** Performance metrics of XGBoost model for 2-class classification problem.

| Confusion Matrix and Statistics |  |  |
| --- | --- | --- |
|  | 2 Class: Wake vs Sleep |  |
|  | Wake | Sleep |
| Wake | 265 | 25 |
| Sleep | 102 | 2908 |
| Statistics by Class: |  |  |
| Sensitivity | 0.72207 |  |
| Specificity | 0.99148 |  |
| Overall Statistics |  |  |
| Accuracy | 0.9615 |  |
| 95% CI | (0.9544,0.9678) |  |
| Kappa | 0.7857 |  |

We also investigated the contribution of each signal in classification performance. To this end, we repeated classification in case each signal was added to the feature vector, and the results are listed in Figure 3,4. As is shown, it can be concluded that all three signals play a pivotal role in sleep staging, and adding new signals can improve classification scores. For XGBoost classifier, when EEG signal was used, we achieved accuracy and kappa: 75.95% and 0.6668, respectively. When PPG’s features was added, the obtained accuracy and kappa raised to: 76.68% and 0.6769, respectively, and after adding the accelerometer’s feature, we achieved accuracy: 81.34± 0.76 and kappa: 0.7388±0.0101.

In order to have a well-fitted model, we need a low bias and low variance model. The learning curve is a way to check bias and variance in supervised learning models. The gap between test and train error shows the model’s variance, and if the model has both high training and test error, it would have a low bias problem. To check the performance of the XGBoost model, we plotted the learning curve of model performance and as shown in Figure 5, the model is neither overfitted nor underfitted. In Table 4, the confusion matrix, Sensitivity, and Specificity rate of this model are given for 5-class classification problem. The sensitivity rate obtained for Wake, NREM1, NREM2, NREM3, and REM is 0.78191%, 0.45312%, 0.8479%, 0.9321%, and 0.6915%, respectively, and the Specificity rate for these classes is 0.98016%, 98842%, 0.8474%, 0.9647%, and 0.9633%, respectively. As the results shows, the model’s performance in terms of the sensitivity and specificity measures is very good.

**Figure 5.**
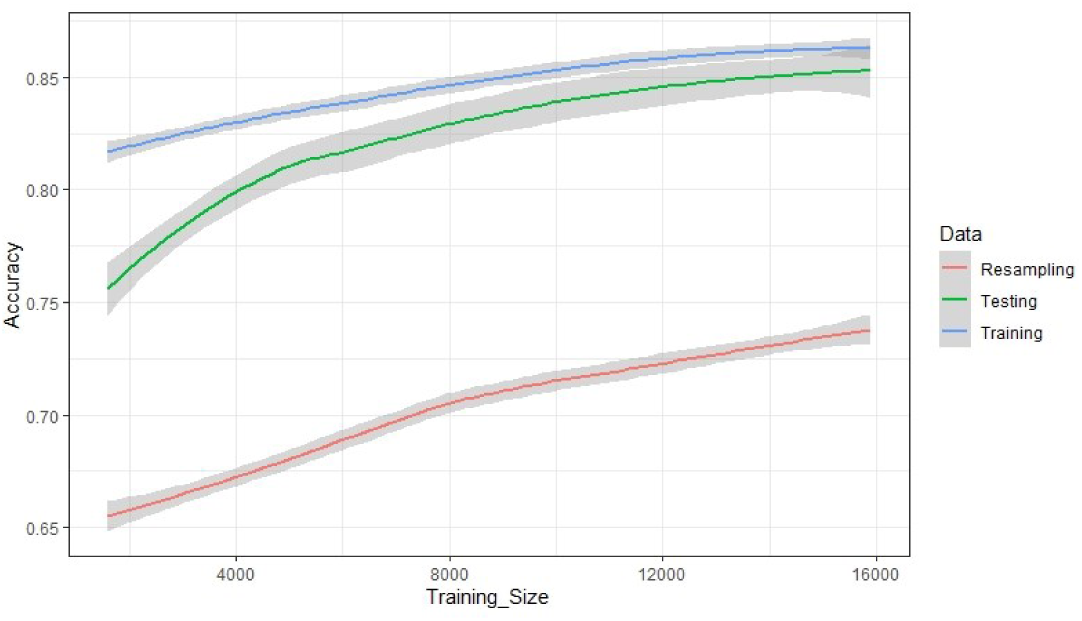
XGBoost model learning curve.

It is worth mentioning that unlike numerous sleep studies that use Pz-Oz channel due to its high classification performance [55], [56], in this study, this channel has not been employed, and instead, we utilized FPZ-O1, FPZ-O2, FPZ-F7, and F7-F8 channels. However, our performance metrics are still promising and can be used in clinical applications. As mentioned before, the objective of automatic sleep stage classification is to propose a model that can be implemented in a device that minimizes discomfort for the subject and performs classification accurately. Considering the application of Dreem headband in sleep analysis and our model performance’s reliability, the proposed model can be used in DH and clinical applications to help sleep experts for sleep disorder diagnosis.

## 4. DISCUSSION

Unfortunately, due to the lack of published papers on this dataset a comparison _on performance ratio_ with others’ works was not possible, which is one of the limitations of our work. We also couldn’t employ other public sleep datasets to evaluate the proposed model since the PSG data in those datasets do not include all three types of signals we used in this study. However, the performance of some of the state-of-the-art sleep stage classification models based on using wearable devices are given and compared in Table 7.

**Table 7.** Comparison of proposed model performance with other state-of-the-art models.

| system | Subject | Dataset | Number of classes | Performance |  |
| --- | --- | --- | --- | --- | --- |
|  |  |  |  | Accuracy | Cohen’s kappa: |
| Beattie et al. [35] | 60 Healthy subject | Accelerometer & PPG | 3class: Wake, Light sleep, REM | 69% | $0.52 \pm 0.14$ |
| Aktaruzzaman et al. [1] | 18 healthy subject | Chest and wrist actigraphy signal | 2 class: Sleep vs wake | 78% for chest signal<br>77% for wrist signal | - |
| Walch et al. [59] | 35 Healthy subjects | acceleration data and heart rate from Apple Watch | 2 class: sleep vs wake<br>3 class: wake, NREM, REM | 90%<br>72% | - |
| Motin et al. [58] | 9393 epochs | wearable fingertip PPG signal | 2 class: Sleep vs wake | 81.10% | - |
| **Our proposed method** | **Healthy and disordered subject**<br>**22596 epochs** | **Dreem Headband** | **2-class: Wake vs Sleep** | **96.15%** | **0.7857** |
|  |  |  | **3class: wake, NREM,REM** | **88.15%** | **0.7154** |
|  |  |  | **5 class** | **81.34%±0.76%** | **0.7388±0.0101** |

In [35], Beattie et al. proposed a model based on using accelerometer and PPG signals acquired by an Apple watch worn by 60 healthy subjects, and their model achieved 69% accuracy for 3-class classification problem and Cohen’s Kappa: 0.52 ± 0.14. In [1] Aktaruzzaman et al. employed chest and wrist actigraphy signal to perform sleep stage classification and evalue their model on 18 healthy subjct’s data. Their model’s accuracy for distinguishing sleep from wake stages was 78% and 77% for chest and wrist signals, respectively. In another study conducted by Walch et al. [57] they recorded acceleration data and heart rate from 35 Healthy subjects worn Apple Watch. Their model’s performance in terms of accuracy for 2-class and 3-class classification problem is 90% and 72%, respectively. In Motin’s work [58], wearable fingertip PPG signal with total 9393 epochs was utilized to classify wake and sleep states. Their model achieved 81.10% accuracy, 81.06% Sensitivity and 82.50% specificity. As mentioned before, although there is a growing interest in using wearable devices for sleep monitoring, one of the limitation of the existing models in this context is that a great number of them can only be used for classifying wake and sleep states and the population consist only healthy subjects. Comparing the performance of our proposed model to other state-of-the-art systems shows that our model not only address the aforementioned problems and outperform existing algorithms, but also can perform 5-class sleep stage classification with a reliable performance. Although the use of different sleep datasets should be taken into consideration when evaluating the performance of these models, this comparison is made to compare the potential of different methods used wearable devices for sleep stage classification.

## 5. CONCLUSION

In this study, we proposed an automatic sleep stage classification model based on using EEG, PPG and accelerometer signals and XGBoost algorithm that can be implemented in wearable devices like Dreem headband and distinguish between different sleep stages accurately. It is worth to highlight some of the significant differences of our work with others:

To the best of our knowledge, except [51] which was conducted to introduce Dreem headband as an alternative to PSG, our work is the first sleep study conducted based on using Dreem headband signals (EEG, PPG, and accelerometer signals). We have also shown the influence of the two latter signals in sleep stage classification. As mentioned before, although using multi-channel signals in sleep studies result in higher performance, this approach is less desirable since recording equipment disrupts patient’s sleep. This study shows that with the proposed model, higher performance can be achieved along with less inconvenience. Furthermore, while in most previous works e.g. Memar et al. [17], Supratak et al. [5], signal acquisition relies on traditional method held in sleep lab which has its own downsides, in this work, polysomnography recordings were acquired using Dreem headband that have several benefits as follow:

➢ It enables signal acquisition at home and eliminates the need for specialists to carry out this task.
➢ It is more convenient for subjects.
➢ It is capable of recording large dataset.
➢ It is cost-effective [51].

Unlike other sleep studies which use less than 16000 epoch for sleep staging [38] we’ve used 22596 epoch in our study that shows our model is robust.

Compared to earlier studies e.g. Boostani et al. [2], Bajaj et al. [26], Berthomier et al. [56], Hsu et al. [60], and Šušmáková et al. [61], in which they mostly use healthy subjects’ data to perform classification, our study included both healthy and sleep disorder subjects; thus, our model can guarantee that the results are reliable for disordered subject, and the system’s performance would not degrade when using the signal acquired from patient having sleep disorder.

## Acknowledgements

This research did not receive any specific grant from funding agencies in the public, commercial, or not-for-profit sectors.

